# Motifs of brain cortical folding from birth to adulthood: structural asymmetry and folding-functional links

**DOI:** 10.1101/2025.03.04.641459

**Authors:** Yourong Guo, Mohamed A. Suliman, Logan Z. J. Williams, Renato Besenczi, Kaili Liang, Simon Dahan, Vanessa Kyriakopoulou, Grainne M. McAlonan, Alexander Hammers, Jonathan O’Muircheartaigh, Emma C. Robinson

## Abstract

Cortical folding of brain is widely regarded as an interplay between genetic programming and biomechanical forces, closely linked to cytoarchitectonic regionalisation. Abnormal folding patterns are frequently observed in neurodevelopmental conditions and psychiatric disorders. However, significant inter-individual variability of secondary and tertiary folds obscures detection of shape biomarkers and confounds investigation of folding-functional relationships. Here we investigate cortical folding heterogeneity at fine scale, using novel hierarchical surface registration (MSM-HT) to parse cortical folding patterns into a representative family of distinct anatomical templates. By applying this technique both to young adults from the Human Connectome Project (HCP) and neonates in the Developing HCP and Brain Imaging in Babies (BIBS) cohorts, we identify and characterize common lobe-wise folding patterns: observing consistency across both age groups, with neonatal samples showing less variation. Crucially, we highlight significant hemispheric asymmetry within the temporal lobe for both adults and neonates, and show that improved correspondence of shape does not translate to improved areal correspondence, affirming previous studies that have pointed to dissociation of folding and functional organisation. This study provides a critical step towards understanding brain asymmetry and complex relationships between folding and function, offering a robust framework to generalise the uncovered cortical folding motifs across datasets and developmental stages.

## Introduction

The human cerebral cortex, with its intricate patterns of sulci and gyri, achieves its remarkable surface area expansion through folding, enabling more complex brain functions. A delicate interplay of genetic factors and biomechanical forces drives this process. Starting at around 11 to 16 weeks, cortical progenitor cells initially establish a ’protomap’ (1); these early cytoarchitectonic areas are genetically defined, largely stable through development, and are closely linked to primary cortical functions.

As cortical development progresses, regional variations in growth, driven by different types of neuronal progenitor cells, along with biomechanical forces, lead to cortical folding (2, 3). Primary folds, which emerge around 20 weeks of gestation, are predominantly shaped by genetic influences, resulting in conserved structures across individuals. Unlike primary folds, secondary and tertiary folds, mostly shaped by week 38, are influenced significantly by intrauterine environmental factors (4) and biomechanical forces, leading to substantial variability and asymmetry. Even with genetically identical individuals (monozygotic twins), significant variability in folding patterns has been observed, highlighting the highly dynamic biomechanical process (5).

It has been challenging to characterise individual variability of cortical folding. In the absence of big open-data collections, and without access to advanced image processing tools, early work was necessarily observational and performed on small samples of brains. For instance, Ono et al. (6) performed a careful examination of 25 brains postmortem, resulting in a detailed description of variabilities of length, branching, connections, and interruptions of major sulci throughout the *entire* brain. Such observational studies are highly time-intensive and dependent on neuroanatomical expertise but do offer detailed characterisation of whole brain cortical shape (7). More recent data-driven, automated methods (8–14) offer less labourintensive alternatives; but so far have been methodologically-constrained to independent investigation of a subset of sulci; limiting scope for exploratory analyses into the extent to which cortical folding patterns co-vary within or between hemispheres, or align with cortical areas.

Though it is often assumed that morphology and function are tightly coupled, this relationship may not hold across the entire brain. While primary cortical folds tend to align with cytoarchitectonic areas supporting sensory/motor functions (15–18), evidence suggests that this relationship is less straightforward in regions supporting complex abilities (19, 20). As pointed out very early by Brodmann, folding landmarks do not correspond to cytoarchitectonic borders precisely (21). For instance, the presence or absence of paracingulate sulci does not affect the existence of its associated cytoarchitectonic areas (area 32) (22).

From a more global perspective of brain function, neuropsychiatric and neurodevelopmental disorders have been associated with atypical cortical shapes (both regionally and across the whole brain) (23–28). However, associating shape with more subtle phenotypes or genotypes has been challenged by marked typical heterogeneity of folding across the population. There is a clear need for analytic approaches that consider this variability to provide a better baseline of what “typical” can be (29). A better analytical framework that addresses heterogeneity in normal anatomy could address the problem of inconsistent and variable findings for these disorders (30, 31).

Disentangling natural cortical shape variability from clinically or cognitively salient features starts with improved characterisation of the scale of cortical shape variability across large populations. To address this, here we propose a novel hierarchical surface-registration approach (Multimodal Surface Matching with Hierarchical Templates, MSM-HT) that clusters - then *aligns* - individual hemispheres into groups that share patterns of cortical folding; resulting in a family of sharp templates, which collectively characterise the most commonly expressed cortical shape motifs, within each of the frontal, temporal and parietal-occiptal lobes. These folding motifs strongly resemble those described in post-mortem studies (6), while benefiting from the scale afforded from big-data analyses to support investigation into the prevalence and asymmetry of motifs. Moreover, propagation of spatial mappings to functional imaging data critically supports investigation into the co-localisation of cortical areas and folds.

We validate our findings in both 1110 young adults, from the Human Connectome Project (HCP), and in 781 term-age neonates from the developing HCP (dHCP) and the Brain Imaging in Babies (BIBS) studies. We reveal consistent yet evolving development of cortical folding from neonates to young adults. At term and adult ages, we found the temporal lobe motifs to be asymmetric, indicating that this morphological asymmetry is already present at birth. Importantly, the folding-function relationship was found not to be tightly correlated in secondary and tertiary-order folds. Altogether, these findings provide a unique perspective on how genetic, structural, and functional processes integrate to shape cortical organization.

## Results

A key assumption underlying most human neuroimaging studies is that - through spatial normalisation - it is possible to map all imaging data to a single population average space, in which common structures (such as sulci and gyri) overlap; this then allows for straightforward comparison and modelling of these features during downstream analyses. Since optimisation of these mappings is ill-posed (with many good solutions) image registration algorithms are typically regularised with diffeomorphic constraints, which impose smooth and invertible transforms that allow no folding or tearing of the deformation grid. This contradicts the body of evidence, described above, which shows that equivalent sulci can appear as continuous in some individuals, but branched, or split, in others. In this paper, we explore a hierarchical image registration framework (MSM-HT), which leverages learning-based methods (32) to perform fast diffeomorphic-alignment between all pairs of hemispheres, from a large cohort; to then intuitively assess how similar cortical folding patterns are, simply in terms of how easily they can be diffeomorphically aligned; this then allows common folding patterns to be clustered out and co-aligned using classic multimodal surface matching (MSM) (33, 34) (see Hierarchical MSM registration (MSM-HT)).

### Characterising modes of cortical shape variation and their frequency

Thirty templates were generated separately for each of the frontal, parietal-occipital and temporal lobes based on HCP young adult data. This was achieved by first assessing the similarity of each pair of lobes from the overlap of their folding patterns, following learning-based alignment; then performing hierarchical clustering, where the resulting dendrograms were thresholded at a fixed value to allow for the assessment of incidence rates for each folding pattern (Supplementary Fig. 1,2 and 3). Aligning data in this way generated a set of templates shown by curvature maps, where larger values represent gyral crown. (Fig. 1 and Supplementary Fig. 1,2 and 3) which show good agreement with the descriptions of Ono’s atlas (6), particularly with respect to segment variations and branching patterns.

**Fig. 1.**
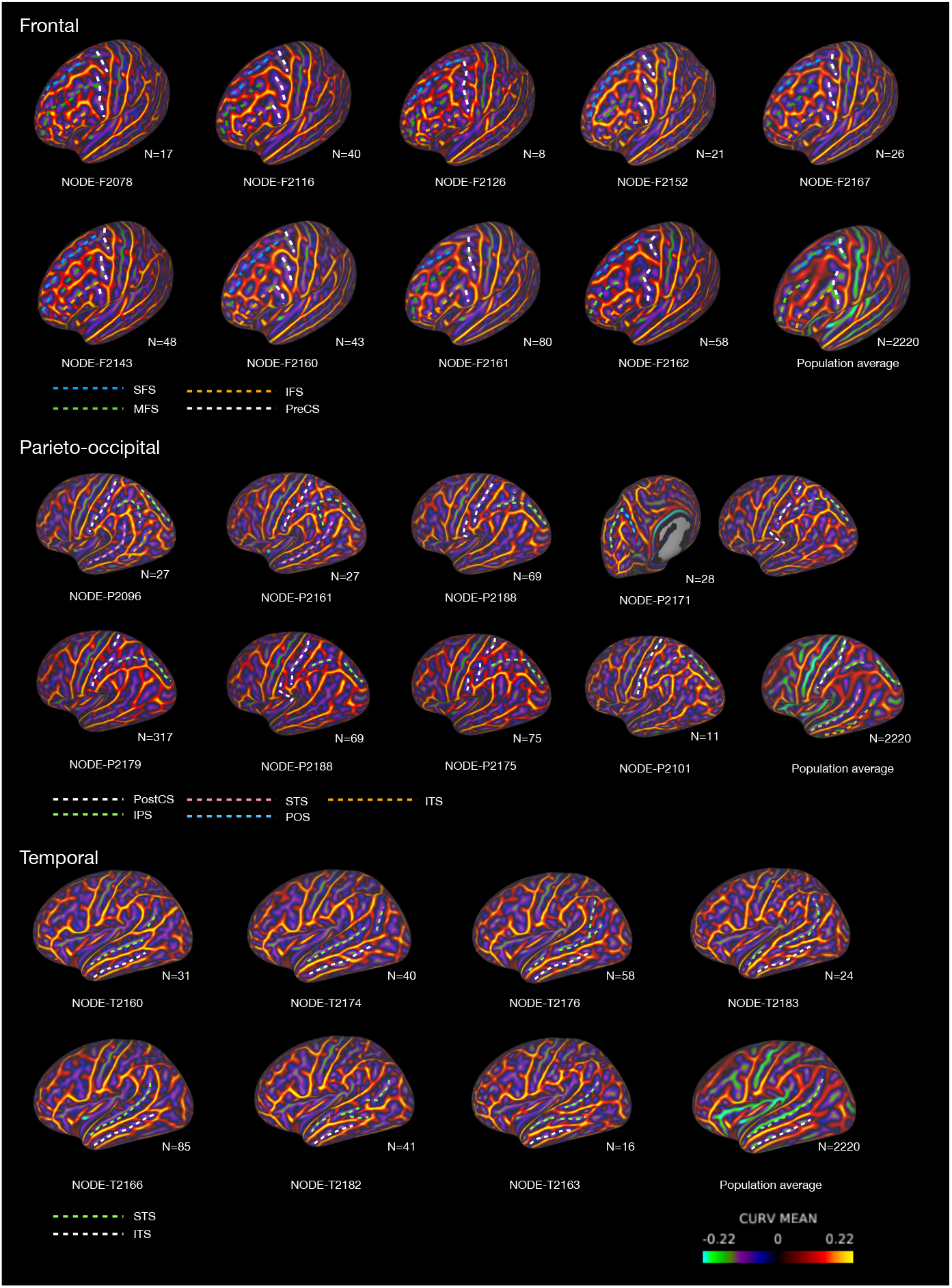
Distinctive common folding motifs through MSM-HT. The cortical folding motifs are shown using curvature, where larger values represent gyral crown. **Top**, representative folding motifs in the frontal lobe. Variability of SFS, IFS, MFS, and PreCS is observed. Cluster F2126 exhibit uninterrupted precentral sulci, while more commonly this sulcus forms 2 or 3 segments, as represented in clusters F2078 and F2160. The PreCS is typically connected to the SFS, with the exception of cluster F2161. MFS shows the most variability, with as many as three or four segments in some clusters (e.g. cluster F2143). A rare connection between the MFS and the anterior segment of IFS is observed in cluster F2078. Connection of the SFS and IFS via MFS segments (e.g., cluster F2078, F2133, F2178) was also identified. **Middle**, representative folding motifs in parieto-occipital lobe. We observed two segments (e.g., clusters P2188 and P2101), three segments (cluster P2175), and uninterrupted PostCS (e.g., cluster P2096); The middle segment of PostCS was found to be connected to the IPS in cluster P2175. Furthermore, in 8 clusters the PostCS was truly connected to the Sylvian fissure (e.g., cluster P2171). The IPS was found to form 1-3 segments, in some cases with ascending side branches (e.g., cluster P2188) or descending side branches (e.g., clusters P2096, P2161). Of these, cluster P2096 represents one example where the descending side branch trends towards the ITS; whereas, in cluster P2161 it connects with the STS. Cluster P2171 represents IPS branches connecting with parietal-occipital sulci (POS). **Bottom**, in the temporal lobe, the STS and ITS display variants with between 1 and 4 segments. The STS has more branching patterns; whereas the ITS displays ’ladder like’ small segments (e.g., cluster T2174). As recorded in Ono’s Atlas, in 77.5% of cases, the STS extends singularly into the AS (e.g., cluster T2166); whereas, in 28.1% of cases (e.g cluster T2176) it displays a double ending - also projecting into the anterior occipital sulcus; with cluster T2169 showing it extends all the way into the occipital lobe. In clusters T2183, T2182, T2163 and T2176 a branched STS connects with the ITS. In cluster T2183 (N=24, 1.1% of the cases), we found a rare pattern in which the STS connected with the Sylvian fissure.

Focusing first on the frontal lobe, the most common pattern presents with two segments of precentral sulcus (PreCS) separated by middle frontal gyrus (MFG); two segments of superior frontal sulcus (SFS); and one or two segments of inferior frontal sulcus (IFS), of which the posterior end connects with the inferior segment of the PreCS. We make the following key observations: 1) uninterrupted precentral sulci (as seen in cluster F2126) are extremely rare, accounting for only 8 hemispheres, and are not described in Ono’s work. More commonly this sulcus forms 2 or 3 segments, as represented in clusters F2078 and F2160. 2) Typically, the PreCS is connected to the SFS, with the exception of cluster F2161 (N=80). 3) In general, the middle frontal sulcus (MFS, termed intermediate frontal sulcus in Ono et al. (6)) shows the most variability, with as many as three or four segments in some clusters (e.g. cluster F2143). In one instance, we observe a rare connection between the MFS and the anterior segment of IFS (cluster F2078, N=17). Connection of the SFS and IFS via MFS segments (e.g., clusters F2078, F2133, F2178) were also identified.

The most common patterns of the parietal-occipital lobe include one or two segments of postcentral sulcus (PostCS); two segments of intraparietal sulcus (IPS); the anterior end of IPS is connected to the PostCS; and the posterior end of IPS is connected to occipital lobe. We found that uninterrupted PostCS are common - accounting for 17 of 30 clusters, of all hemispheres (e.g., cluster P2096); two segments are also frequent (12 of 30 clusters. e.g., clusters P2188 and P2101) but three segments are uncommon: accounting for 1 of 30 clusters (roughly 3.4% of all hemispheres), and its middle segment was found to be connected to the IPS (cluster P2175). Furthermore, in 8 clusters the PostCS was truly connected to the Sylvian fissure (e.g., cluster P2171). Considering instead the IPS, this was found to form 1-3 segments, in some cases with ascending side branches (e.g., cluster P2188) or descending side branches (e.g., clusters P2096, P2161). Of these, cluster P2096 represents one example where the descending side branch trends towards the inferior temporal sulcus (ITS); whereas, in cluster P2161 it connects with the superior temporal sulcus (STS). Cluster P2171 represents IPS branches connecting with parietal-occipital sulci.

In the temporal lobe, the most common folding patterns include one or two segments of STS; three or four segments of ITS; the anterior end of STS locate at temporal pole; the posterior end of STS extent towards parietal lobe angular sulcus (AS). We found that the STS and ITS again display variants with between 1 and 4 segments. The STS has more branching patterns; whereas the ITS displays ‘ladder like’ small segments (e.g., cluster T2174). As recorded in Ono’s Atlas, in 77.5% of cases, the STS extends singularly into the AS (e.g., cluster T2166); whereas, in 28.1% of cases (e.g cluster T2176) it displays a double ending - also projecting into the anterior occipital sulcus; with cluster T2169 (N=31) showing it extends all the way into the occipital lobe. In clusters T2183, T2182, T2163 and T2176 a branched STS connects with the ITS. In cluster T2183 (N=24, 1.1% of the cases), we found an uncommon pattern in which the STS connected with the Sylvian fissure.

We also investigated whether individuals who share the same variant of cortical folding in one lobe, also show similar patterns across other lobes. In total 14 pairs of co-occurrence (12 parietal-temporal and 2 frontal-temporal) achieve corrected-p <0.001 and *κ* > 0.4, indicating moderate to substantial agreement (see Supplementary Results and Supplementary Discussion; however, these are largely characterised by larger cluster sizes and more common folding patterns (see Supplementary Fig. 4)

### Cortical folding motifs are heritable

Heritability can provide some insights into potential genetic mechanisms underpinning cortical folding. Research has shown that early-formed primary folds are heritable, but even monozygotic (MZ) twins can have substantially different folding patterns. We used the shared genes between MZ twins and dizygotic (DZ) twins to investigate the genetic basis of cortical folding. We tested for a difference in frequency of motif co-occurrence in MZ and DZ twins from the young adult HCP dataset using a Chi-squared test, demonstrating that, in all lobar subdivisions, motifs co-occurred more frequently in MZ siblings than DZ. The OR was consistently larger than 1, indicating that MZ twins are more likely to be in the same cluster. (values: *χ*^2^*/*corrected *p/*OR, left frontal: 3.94/0.047/2.16; right frontal: 7.14/0.014/2.99; left parieto-occipital: 10.45/0.004/2.97; right parieto-occipital: 6.75/0.014/2.63; left temporal: 4.51/0.040/2.39; right temporal: 10.45/0.004/2.97).

### Greater asymmetry of the temporal lobe

It is well known that the cortex displays morphological asymmetries, with inter-hemispheric differences in sulcal depth, curvature and thickness that have been linked to functional specialisations such as language, visuospatial processing and handedness (35, 36). The very first structural asymmetry was reported by Geschwind and Levitsky (37), showing that the left planum temporale was larger than the right one and supposed that this feature was later associated with language functions including comprehension and processing. Leroy et al. (38) and Le Guen et al. (39) reported human-specific, right-deeper-than-left sulcal depth in STS and more interruptions in cortical folding in left STS than right, which is likely to be constrained by genes and related to linguistic networks (40). Since our templates were generated through joint clustering of left and right hemispheres, this enabled us to posit the question of whether certain cortical patterns were more left or right-dominated - simply by comparing the frequency of left hemispheres used to build each template. This demonstrated significantly greater asymmetry of the temporal lobe (Fig. 2) relative to the frontal (corrected p = 0.014) and parieto-occipital regions (corrected p = 0.009), with more templates showing leftward bias than rightward bias based on the frequency.

**Fig. 2.**
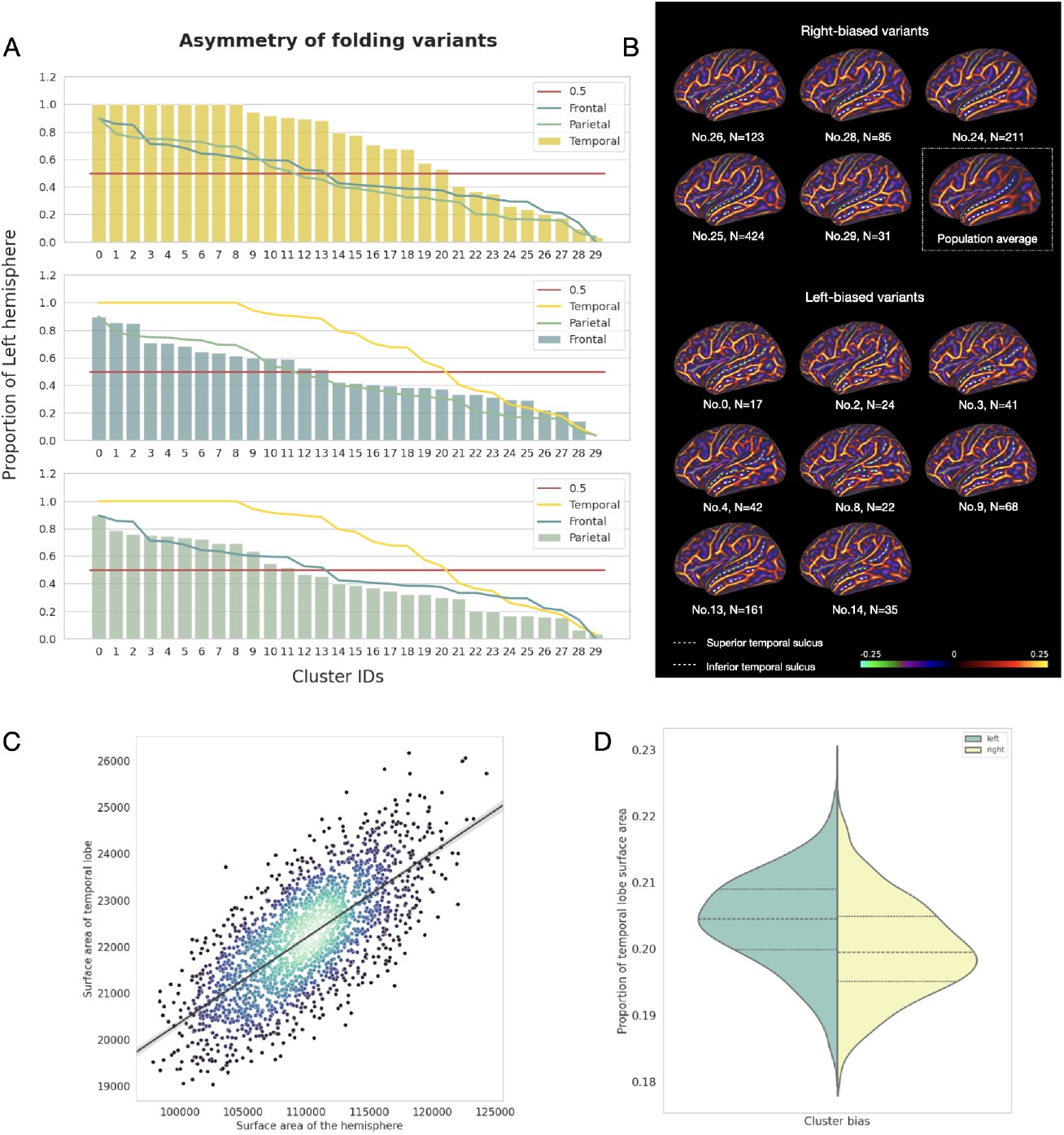
Asymmetry of the temporal lobe cortical folding patterns. **A**. Proportions of left hemispheres used to construct each template, sorted in descending order. There is significantly greater asymmetry of the temporal lobe relative to the frontal (corrected p = 0.014) and parieto-occipital regions (corrected p = 0.009), with more templates showing leftward bias than rightward bias. **B**. Leftward- and rightward-asymmetric templates in temporal lobe. Leftward-asymmetric templates exhibited more atypical folding variation than rightward, clustering into smaller and more variable groups, that display increased interruptions and branches of the STS; whereas rightward-asymmetric templates displayed a more continuous STS. Specifically, No. 2, 3, 4, and 8 showed branches of the middle segments, extending into ITS, while No. 4 also displayed branches of the posterior segments of the STS; **C**. Normalized surface area of the temporal lobe for leftward- and rightward-asymmetric clusters. The surface area of temporal lobe increases with surface area of the entire hemisphere. **D**. The normalised temporal lobe surface area was compared, where leftward-asymmetric folding variants were significantly larger than those of rightward variants (p < 0.001).

Closer examination showed that leftward-asymmetric templates exhibited more atypical folding variation than rightward, clustering into smaller and more variable groups, that display increased interruptions and branches of the STS; whereas rightward-asymmetric templates displayed a more continuous STS. Specifically, No. 2, 3, 4, and 8 in Fig. 2 B showed branches of the middle segments, extending into ITS, while No. 4 also displayed branches of the posterior segments of the STS (Fig. 2 B); This was confirmed by comparing the normalized surface area of this region, showing that leftward-asymmetric folding variants were significantly larger than those of rightward variants (Fig. 2 C, p < 0.001). Both left and rightward-asymmetric variants showed multiple interruptions of the ITS.

### Pattern and asymmetry replicate in neonates

To evaluate how stable these patterns are, across datasets and across development stages, we replicated MSM-HT on term-age scans of neonates from the dHCP and BIBS cohorts. We hypothesised that MSM-HT would identify consistent cortical folding patterns in neonatal data, similar to those found in the HCP dataset, indicating that the framework is robust across different development stages. We calculated the Pearson correlation similarity of mean curvature maps between all neonatal and adult folding variants. For each neonate’s folding variant, the correlation similarities with all HCP folding variants were ranked from the highest to the lowest. In most cases, this indicated strong pairings between specific neonatal and adult counterparts; often with more than one adult template matching to each neonatal template (Fig. 3 A). To illustrate this, the top two most similar adult frontal variants in frontal lobe are shown in Fig. 3 B, with the first folding variants showing strong visual resemblance in terms of special folding characteristics (Top 1 column), marked by the white dash line. For instance, neonatal cluster NEO-F1475 and HCP cluster F2139 both have small gyri interrupting the IFS and separating the MFS from IFS; neonatal cluster NEO-F1491 and HCP cluster F2161 both have middle frontal gyrus (MFG) detached from the precentral gyrus (PreSG); neonatal cluster NEO-F1511 and HCP cluster F2133 both have PreCS connected to MFS; neonatal cluster NEO-F1524 and HCP cluster F2184 both had parallel branches of MFG.

**Fig. 3.**
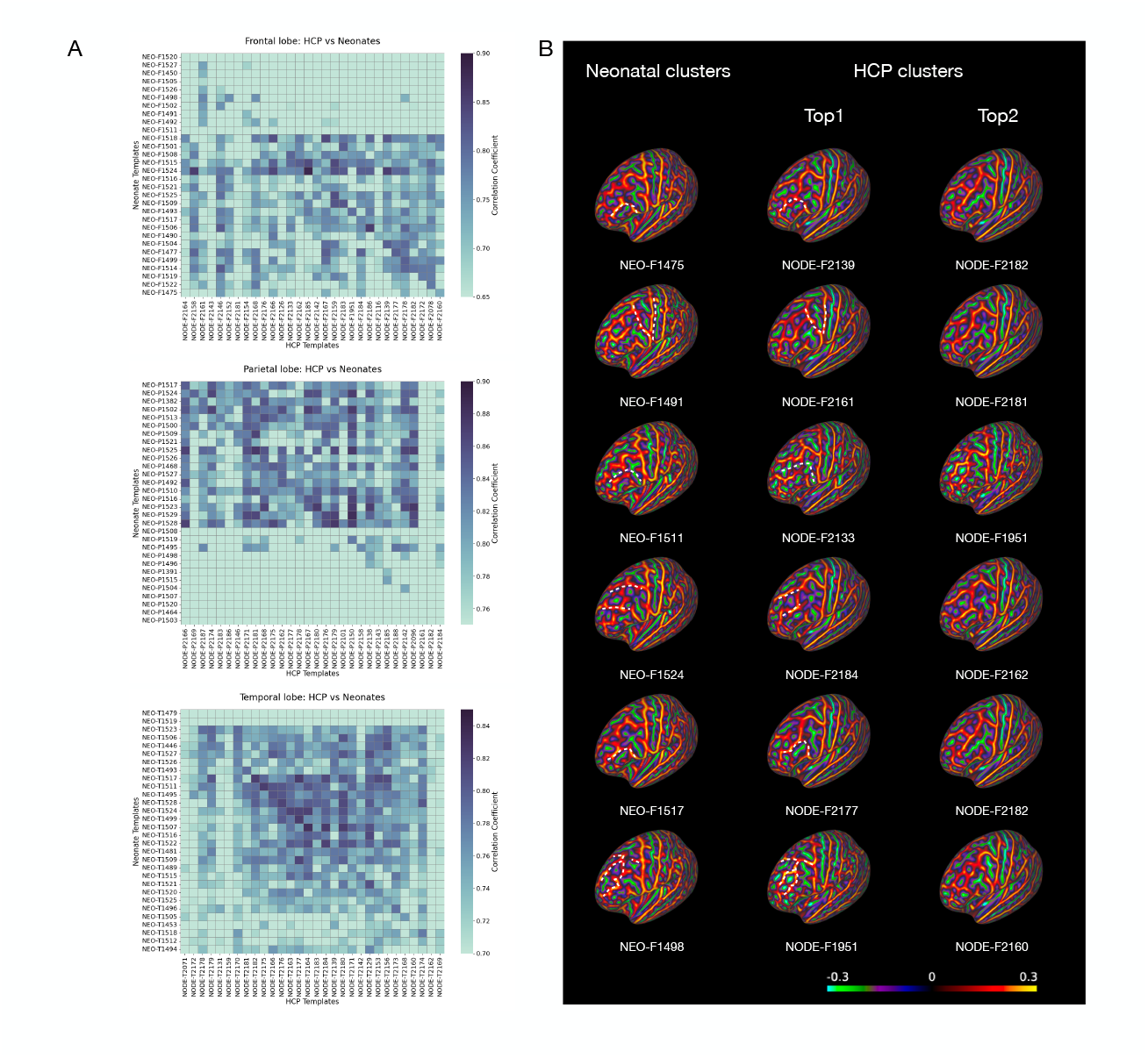
Replication of MSM-HT in the neonatal brain. **A**. Correlation similarity between HCP and neonatal folding pattern. The order of the neonates’ folding patterns was arranged to show the most similar neonatal variants close to the diagonal. In almost all pairings we find a highly correlated match. **B**. Top matches of neonatal and adult folding variants. The first column shows the example frontal folding variants from neonates. The last two columns show the first and second most similar adult folding variants. The Top 1 column folding variants show consistent visual resemblance to neonates in terms of special folding characteristics, and the Top 2 column variants showed far less pronounced visual similarities. Similar patterns were pointed out by the white dash line.

We extended the analysis of cortical folding variants to investigate temporal lobe asymmetry in neonates. While a trend of greater temporal lobe asymmetry compared to other lobes was observed, it did not reach statistical significance (Fig. 4 A, Mann-Whitney U test followed by FDR correction, temporal vs. frontal p = 0.44, corrected p = 0.62; temporal vs. parietal = 0.62, corrected p = 0.62). This might reflect differences in the developmental trajectory of hemispheric folding. Similar to the adult dataset, the leftward asymmetric folding patterns exhibited more interruptions on STS; while the rightward asymmetric folding patterns tended to have continuous STS (Fig. 4 B). Both left and rightward-asymmetric variants showed multiple interruptions of the ITS. Contrary to the adults, the normalized surface area of the temporal lobe showed that leftward-asymmetric folding variants were significantly smaller than those of rightward variants (Fig. 4 C, p < 0.0001). This inverse asymmetry pattern between adults and neonates could suggest that the left hemisphere, particularly the temporal lobe, is still undergoing development postnatally, potentially due to environmental influence and language development afterbirth (41, 42).

**Fig. 4.**
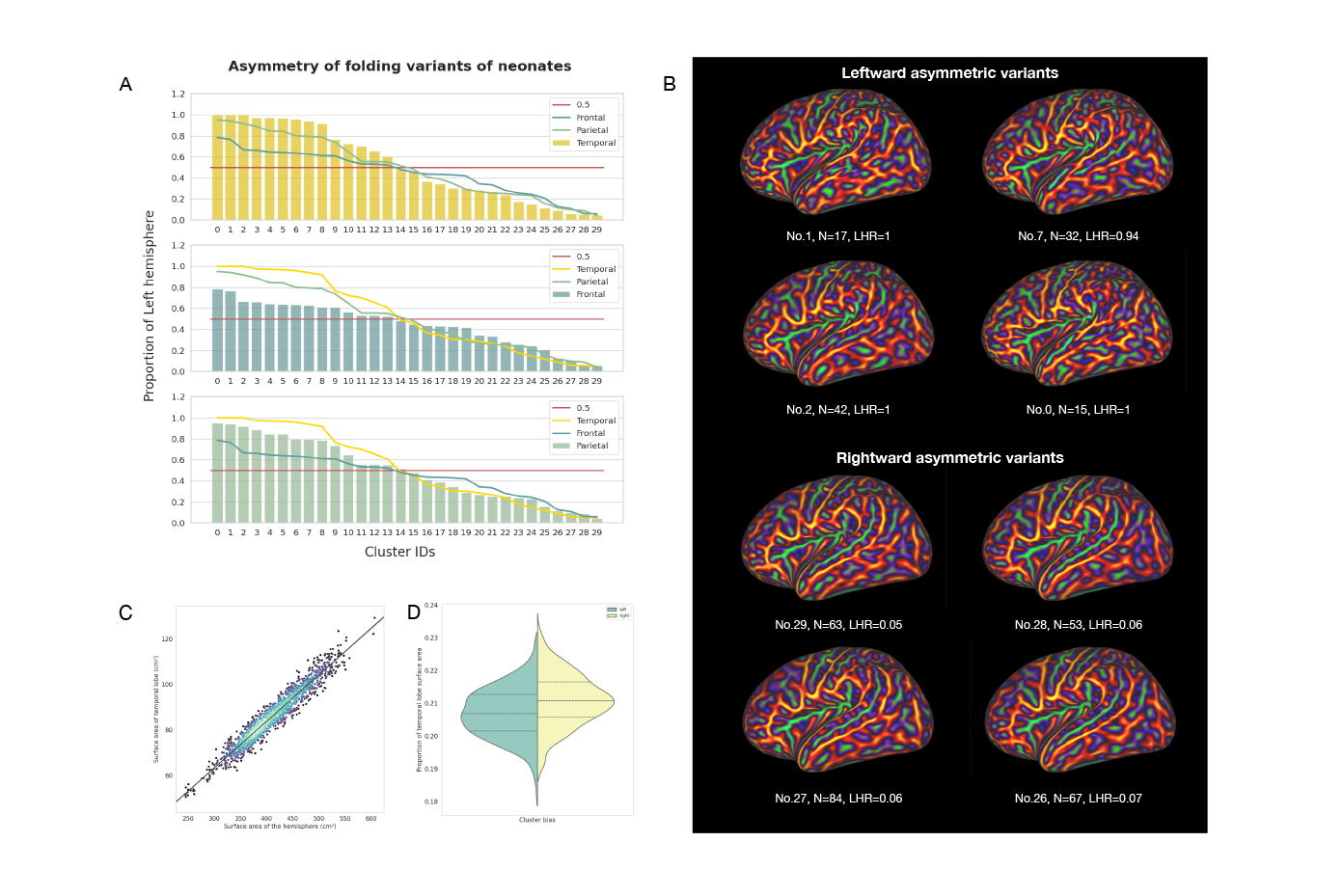
Temporal lobe asymmetry of neonatal datasets. **A**. Proportions of left hemispheres used to construct each template, sorted in descending order. While a trend of greater temporal lobe asymmetry compared to other lobes was observed, it did not reach statistical significance (Mann-Whitney U test followed by FDR correction, temporal vs. frontal p = 0.44, corrected p = 0.62; temporal vs. parietal = 0.62, corrected p = 0.62). **B**. Leftward- and rightward-asymmetric templates in temporal lobe. The leftward asymmetric folding patterns exhibited more interruptions on STS; while the rightward asymmetric folding patterns tended to have continuous STS. Both left and rightward-asymmetric variants showed multiple interruptions of the ITS. The color bar is the same as Fig. 3. **C**. Normalized surface area of the temporal lobe for leftward- and rightward-asymmetric clusters. Similarly to the adults, the surface area of temporal lobe increased with the surface area of the entire hemisphere. **D**. Contrary to the adults, the normalized surface area of the temporal lobe shows that leftward-asymmetric folding variants were significantly smaller than those of rightward variants (p < 0.0001).

### Functional areas lack spatial correspondence with folding patterns

Development of cortical folding has long been thought to relate to the development of functional areas. It was hypothesised that the cortical function regionalisation happens first, then cortical folding is induced related to the spatial arrangement of these functional areas (1, 43). As shown in Fig. 5, the current work has proved that MSM-HT significantly improves the alignment of structural features including sulcal depth and curvature relative to the classical approach of aligning all images directly to a single population template (i.e., MSM-P, the template used here corresponds to the official MSMSulc template used in the HCP pipelines). The mean sulcal depth and curvature maps were sharper and revealed more details of the folding pattern. Accordingly, if the folding-functional relationship is strong, it is reasonable to assume that MSM-HT would also improve the overlap of functional activation patterns. This was therefore tested by projecting HCP task and resting-state fMRI into the MSM-HT and MSM-P template spaces.

**Fig. 5.**
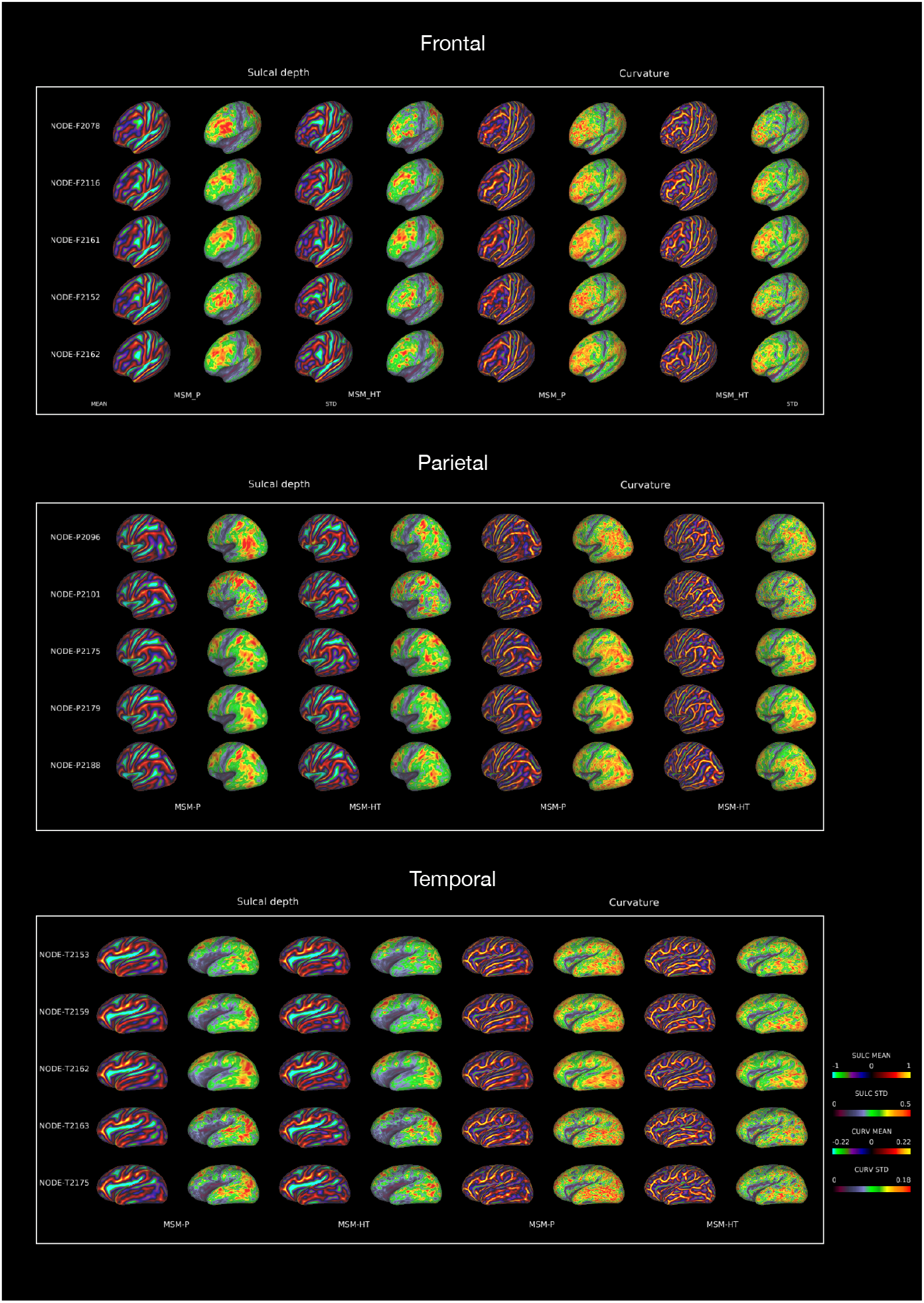
Mean and STD map of example cortical folding clusters using MSM-P and MSM-HT. **Top**, frontal example clusters. **Middle**, parieto-occipital example clusters. **Bottom**, temporal example clusters. The mean and STD maps of sulcal depth and curvature are compared between MSM-P and MSM-HT: column 1, 2, 3 & 4 are mean and STD maps of sulcal depth, registered with MSM-P and MSM-HT respectively; column 5, 6, 7 & 8 are mean and STD maps of curvature, registered with MSM-P and MSM-HT respectively. The color bars are adjusted to be the same for each of sulcal depth mean, STD, curvature mean, STD for a fair comparison between MSM-P and MSM-HT.

Using the HCP task-based fMRI (tfMRI) data, we aligned language, motor, social, and working memory contrasts with the two different registration methods. The area distortion of both MSM-HT and MSM-P was controlled to be equivalent (Supplementary Fig. 6), in order to exclude the possibility that the strength of the shape-driven surface registration might influence functional overlap. The resulting within-cluster group average activation z-stats maps show that functional activations were either similarly localised in both MSM-HT or MSM-P e.g. for the language-math contrast of the frontal lobe cluster F2161 (Fig. 6A), or in some cases were slightly less well-aligned for MSM-HT. To quantify this effect, we evaluated functional alignment using *cluster mass*: a measure of the strength and spatial concentration of significant activation vertices for which higher values indicate better overlap of task activations across the group (Fig. 6B). Cluster mass was significantly lower in MSM-HT than MSM-P for frontal and parietal lobes in all four tasks (corrected *p <* 0.01). However, no significant differences were observed for the temporal lobe in any tasks. This suggests that MSM-HT leads to a less concentrated spatial alignment of functional activation in frontal and parietal lobes, while temporal lobe is not affected. This could be explained by the fact that the placement of architectonic areas is not constrained by morphological landmarks (sulci and gyri) (21).

**Fig. 6.**
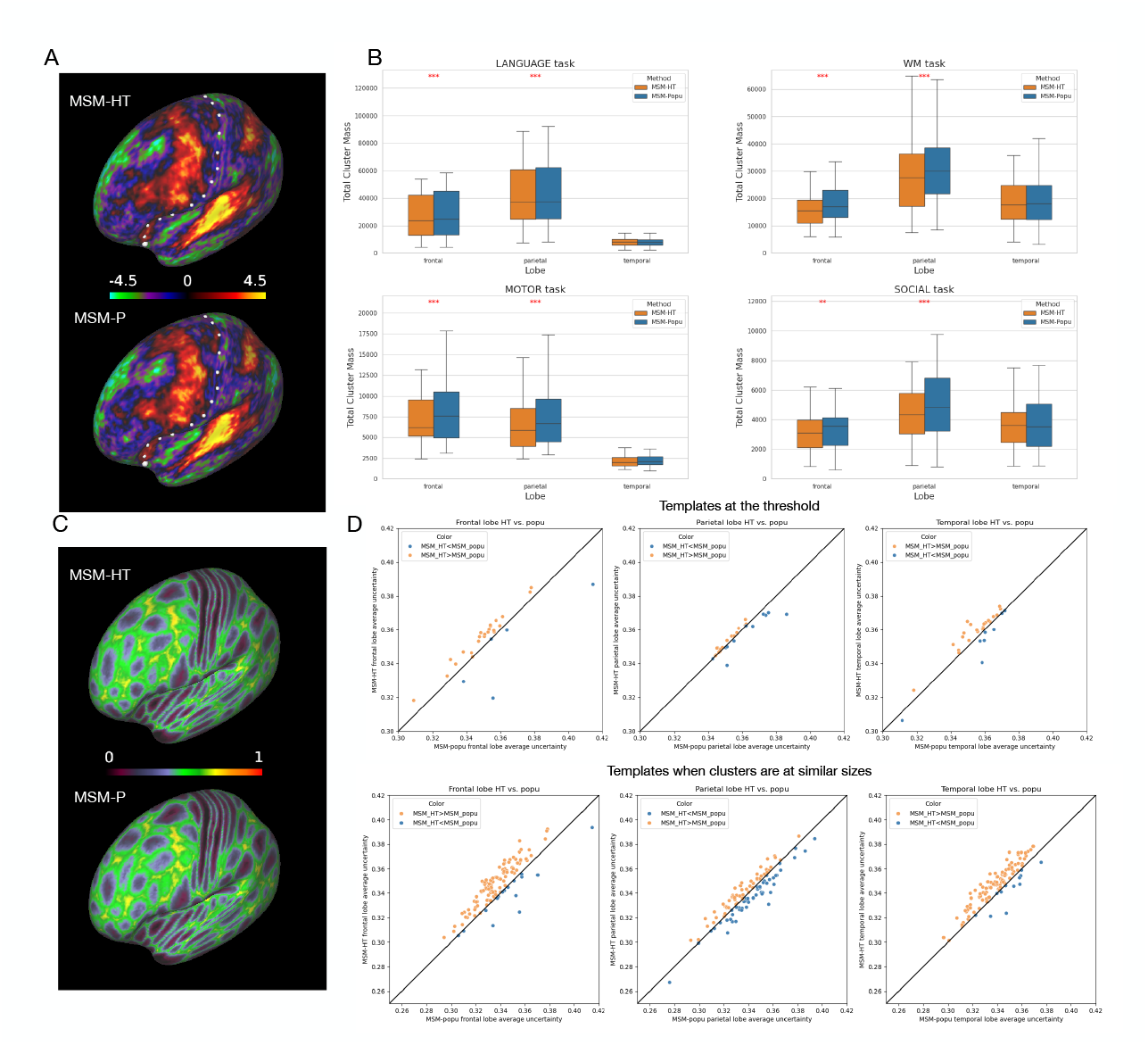
Folding-functional relationship. **A**. Example cluster-average functional activation of language-math contrast using MSM-HT and MSM-P (frontal lobe cluster F2161). Functional activations were similarly localised and concentrated using both MSM-HT and MSM-P registration methods **B**. Cluster mass distribution across three lobes for language, working memory, motor, and social tasks. The cluster mass was significantly different between MSM-HT and MSM-P for frontal and parietal lobes in all four tasks (corrected *p <* 0.01), where cluster mass is higher in MSM-P alignment. However, no significant differences were observed for the temporal lobe in any tasks. **C**. Example multimodal parcellation cluster uncertainty using MSM-HT and MSM-P (frontal lobe cluster F2161). **D**. A scatter plot comparing within-lobe average areal uncertainty between MSM-HT and MSM-P. The top row showed average areal uncertainty for folding motifs generated using a threshold that returned 30 clusters. The bottom row showed average areal uncertainty for folding motifs generated in comparable sizes. See Folding-function relationship.

To investigate how MSM-HT affects general functional alignment beyond the four tasks considered, we used the spatial mappings from both MSM-HT and MSM-P to align individual HCP multimodal cortical areal parcellations (44). These parcellations, driven by a broader set of functional features such as resting-state spatial maps, T1w/T2w myelin maps and cortical thickness, provide a more comprehensive view of functional organisation than specific tasks. Fig. 6C shows the areal uncertainty of the multimodal parcellation, of example cluster F2161, for both MSM-HT and MSM-P. Average areal uncertainty is similar (Fig. 6D), which reinforces the tfMRI finding that folding and function are not related as tightly as thought. Instead, other factors might drive functional alignment, such as cortical connectivity and plasticity.

## Discussion

Patterns of cortical folding result from the long process of structural and functional cortical development and offer a window into misdevelopment. Our study was motivated by a need to uncover a comprehensive baseline of “typical” cortical folding, to increase our sensitivity to detect morphological motifs of behaviour and disease. To achieve this, we parsed heterogeneous patterns of cortical folding into a discrete set of typical templates for each anatomical lobe. Our motifs aligned with variants manually defined by Ono et al. (6), and our results uncovered both common cortical variants (i.e., seen in at least 5% of the sample) and known, but rarer, variants (such as the single-segment PreCS in cluster F2126). This anatomical foundation provides a novel framework for exploring structural brain organisation at a larger scale.

Our motifs, identified in adults, are representative of the earliest stages of post-natal life. The close resemblance between the identified neonatal and adult folding motifs suggests the developmental consistency of major cortical folding patterns established prenatally or shortly after birth across different life stages. However, there is evidence that temporal lobe folds are still under fast development at this stage (45). For instance, comparing term-age neonates and adults, Hill et al. (41) reported that the lateral temporal and anterior cingulate folding can still be curving and branching after birth. In our result, while some folding patterns in neonates mirrored those found in adults, the diversity of these patterns was notably less in the neonatal brain, especially for frontal and temporal lobe.

Our study shows the left temporal lobe exhibited more folding variability than the right: the major difference lay in STS, where the leftward-symmetric folding variants have more branching patterns and interruptions by ’plis de passage’. This asymmetry trend has been observed both in neonates and adults. The increased structural complexity on the left may serve as a structural basis for the lateralization of language processing, as supported by Bodin et al. (46), who found dense bundles of U-shaped fibres related to the positions of the plis de passage. In line with this, we discovered significantly higher normalized temporal lobe surface areas in the left hemisphere compared to the right in adult data, indicating more computation related to language processing.

Notably, temporal lobes are differentially asymmetric at the two age points. In the early developmental stage, the right temporal lobe matures earlier than the left in cortical folding complexity (42). Our finding in neonatal datasets supported this with less pronounced temporal lobe asymmetry relative to other lobes, with fewer leftward-asymmetric temporal lobe templates relative to adults. Moreover, rightward-asymmetric folding clusters were found to have higher temporal lobe surface area than the leftward-asymmetric clusters, where this was reversed in adults. This aligns with findings from Williams et al. (47), who found significantly higher leftward surface area asymmetry in adults compared to neonates along the STS and STG. Moreover, it is known that the right temporal lobe specialises in facial recognition and social cognition (38); fundamental to the development of social bonding with parents; and its structure is more genetically controlled and well-established at birth than left temporal lobe (48). On the other hand, the left temporal lobe is specialised for language processing (38), which is shaped and influenced by the functional stimuli after birth, such as language exposure (42). Roughly two years after birth, the synaptic pruning starts (49) and facial discrimination is not as important a function in this stage. We hypothesise this process gradually changes the relative surface area of left and right temporal lobes, such that by the time that cortical folds approach stability, the temporal lobe presents more pronounced asymmetry.

It therefore seems that, from the link between structural asymmetry and functional lateralisation, the brain cortical folding development is related to functional development. Indeed, theories have been proposed that suggest that early placement of cortical folds might relate to the formation of functional regions (1, 43). However, this folding-functional link hypothesis has been validated mostly in primary functional areas, including the primary motor and visual cortex (19); whereas, it has become clear from our study and others (20, 41) that the folding-function relationship in higher-order areas is more complicated. In these areas, folding is much more irregular and the association with function is much more dissociated. The reason why cortical folding localises more tightly with primary functional regions but seems stochastic in association functional areas might be explained by several factors. In line with previous heritability research in HCP (4), our study found a stable genetic influence within lobes in the adult brain, indicated by MZ twins being significantly more likely to fall into the same folding clusters than the DZ twins. Indeed, cortical areas are genetically programmed with specific functional regions, where early cortical folds are formed related to them (1, 43, 50). Furthermore, recent studies show that gene expression is highly localised prenatally and this persists towards adulthood (51). The genetic gradient also sharply changes across the first-forming sulci (52). Altogether, these studies suggest that genetic control might explain why the first-forming sulci along the time course exhibits less between-individual variability than later-forming sulci. However, later in the folding process, the competition between genetic control and biomechanical pressures might start to bias towards the latter since gene transcription becomes more synchronised across regions in the late prenatal stage (53). The balance between different biomechanical forces, such as local cell proliferation and white matter tract tethering, can be highly sensitive (3); small differences in the earlier developmental stage (e.g., variations in proliferation) and later neuron positioning through migration can result in significant changes in later-forming folds, leading to highly irregular sulci (52, 54). On the other hand, while overall brain shape (primary and secondary folds) is mostly fixed at birth, functional development continues into adulthood; therefore functional plasticity may be another factor in the final placement of functional areas. The early positioning can be regulated by signalling molecules discharged from multiple patterning centres (55). Moreover, multiple environmental factors such as social status, language exposure, and pathological circumstances can influence the development of association networks and higher-order functions throughout childhood and adulthood (56–58).

MSM-HT represents a powerful tool for uncovering and characterizing cortical folding patterns. By emphasizing the neuroscience findings, this study contributes to our understanding of the complex relationships between brain structure, function, and development, paving the way for future research that can build on these insights. However, there are several limitations to consider. First, the comparison to Ono’s Atlas should be interpreted with caution. The lobe-wise clustering approach provides a broad view but lacks the resolution to identify variability within individual sulci, which is detailed in a small-scale postmortem study. This methodological difference presents challenges for direct comparison. Second, when comparing the MZ twins or DZ twins, in some instances twins may not fall into the same clusters but similar clusters. This results in their cortical folding pattern being treated as distinct in statistical analysis, introducing uncertainty and potentially blurring the statistical result. Finally, it is difficult to perform tfMRI on neonates, which lack a direct comparison to our result of adult tfMRI. Moreover, the low spatial resolution of fMRI relative to the size of the neonatal brain also results in partial-volume effect, which makes it more challenging to identify some of the components observed in the adults (59). The diversity of populations and age groups has long confounded the search for shape biomarkers of neurological disorders, which limited the scope of biomarkers for neurological disorders mostly in primary folds (24, 26). Therefore, for future work, we aim to expand the application of MSM-HT to a developmental perspective to enhance the generalisability of the findings and identify population-specific patterns of cortical development. Additional data types, such as functional connectivity or genetic information, can also be integrated to provide a more comprehensive understanding of cortical folding. Such efforts will be critical for disentangling the disease-specific patterns from population variability to facilitate biomarker discovery.

## Method

### Datasets

The data used for the development and validation of the proposed method correspond to structural and functional imaging data from 2220 cortical hemispheres derived from the young adult HCP. Cortical folding templates were subsequently replicated in a neonatal cohort (1562 hemispheres) derived from the dHCP and BIBS.

#### HCP Young Adult

Data were collected from 1110 young adults aged 28.8 *±* 3.7 years (604 biologically females) on a Siemens Skyra “Connectom” scanner, with a standard Siemens 32-channel head coil (60). T1w scans were acquired with a 3D MPRAGE sequence (TR=2400 ms, TE=2.14 ms, TI=1000ms, voxel size=0.7 mm isotropic); whereas, T2w scans were acquired with a 3D T2-SPACE sequence (TR=3200, TE=565, voxel size=0.7 mm isotropic). For functional acquisition, resting-state fMRI was acquired with a Gradient-echo EPI sequence (TR=720ms, TE=33.1ms, flip angle=52 degrees, multiband factor 8, 60 minutes for 4 runs, voxel size=2mm isotropic); tfMRI was acquired with the same sequence parameters as resting-state fMRI (7 tasks including Language, Working Memory, Motor, Social Cognition, Gambling, Emotion Processing, Relational Processing, 48 minutes total). Full details regarding acquisition and participant inclusion criteria are described in (61). All data is open and accessed through ConnectomeDB.

#### dHCP and BIBS

dHCP and BIBS are parallel studies run at the Evelina London Children’s Hospital, St Thomas, London, that dual-consented participants; dHCP was a community cohort which included preterm born children and BIBS recruited neonates with and without familial history of neurodevelopmental and/or psychiatric conditions such as autism, attention deficit hyperactivity disorder (ADHD), or mood disorder. Full inclusion criteria for dHCP and BIBS control subjects are described in (62). All datasets were acquired on a Philips Achieva 3T scanner at the Evelina Newborn Imaging Centre with a dedicated 32-channel neonatal head coil (62, 63). 781 neonates were used, of which 648 were term-born (scan age 41.5 *±* 1.7 weeks, 297 biologically female); 133 were preterm-born (scan age: 41.3 *±* 1.7 weeks, 59 biologically female); 44 had a known family history of autism; 22 had a know family history of ADHD; 10 had a mother with mood disorder. Structural T2w MRI was acquired with a TSE sequence: TR=12000 ms; TE=156 ms; SENSE factors: 2.27 axial and 2.66 sagittal; in-plane resolution 0.8 *×* 0.8, slice thickness: 1.6 mm, and slice overlap: 0.8 mm.

### Image Pre-processing

#### HCP

Adult data were processed using an adapted version of the FreeSurfer pipeline (64), described in (61). This involved tissue segmentation and extraction of inner (white) and outer (pial) cortical surfaces, inflation (for improved visualisation of sulci), and projection to a sphere. The white and pial surfaces define the cortical ribbon. Integration over the energy functional used for inflation generates sulcal depth maps, which summarise coarse-scale folding; we also used mean curvature (estimated from the average of two principal curvatures of the white surface) and cortical label maps corresponding to the 34 regions of the Desikan Killiany atlas (65). For functional data, the timeseries first underwent motion correction, and distortion correction; then the voxels within the cortical ribbon were mapped to the cortical surface, transformed to the 32k_FS_LR mesh with surface registration, and smoothed within the surface with a 2mm FWHM Gaussian kernel. The task contrasts were estimated within each run then each subject using general linear model (GLM) analyses implemented in FEAT (66, 67).

#### dHCP and BIBS

Neonatal surfaces were extracted using a recently published bespoke deep learning-based pipeline (68), trained on the outputs from the original dHCP pipeline (69). This pipeline adapts the FreeSurfer pipeline to the very different MRI properties of neonatal scans, which have inverted tissue contrast and relatively low resolution. Visual quality control (QC) of the surface output from the learning-based pipeline was improved in terms of anatomical and topological correctness (68). We use neonatal spheres, sulcal depth maps, mean curvature, and cortical labels from the DRAW-EM regional atlas (70–72), which differs from the Desikan-Killiany atlas, but ensures accurate segmentation of the neonatal brain structures. For more details about the deep learning-based neonatal pipeline, see Supplementary Note Deep learning-based neonatal pipeline.

### Hierarchical MSM registration (MSM-HT)

#### Overview

MSM-HT leverages the complementary benefits of two separate tools for cortical surface registration: classic MSM (33, 34, 73) and its learning-based implementation (Deep-Discrete spherical Registration, DDR) (32, 74). Both methods solve the same objective: deforming an input (or moving) mesh until its features (e.g., sulci and gyri) better overlap with those of a reference surface whilst penalising extreme (biologically implausible) deformations; however, while MSM offers theoretical guarantees over the smoothness and invertibility of its deformation fields, DDR offers fast approximate solutions that are better suited to performing pairwise alignment between all cortical hemispheres, in large cohorts. For this reason, we used DDR to coarsely align large numbers of pairs of hemispheres and MSM to generate the final templates. For more details on each of these methods, see (32–34, 74–76).

The resulting framework of MSM-HT is shown in Fig. 7. Starting with the null hypothesis that there is no asymmetry of the cerebral cortex, we first mirror-flip all right hemispheres (into rigid alignment with left hemispheres) using Workbench Command; then co-register all pairs of sulcal depth maps using DDR. Once sulcal depth maps are coarsely aligned, the overlap of cortical folding patterns for each pair of hemispheres may be separately assessed for each of the frontal, parieto-occipital, and temporal lobes - using a combination of Dice overlap and cross-correlation. Agglomerative hierarchical clustering is then applied to each of the resulting similarity matrices, and the hierarchy is thresholded at the similarity score that returns thirty clusters for each lobe. MSM is then used to co-register curvature maps of individuals within each cluster to generate a template for each cluster - that summarises the shape of folds within that cluster. Finally, all templates are further registered through the hierarchical path to a common space.

**Fig. 7.**
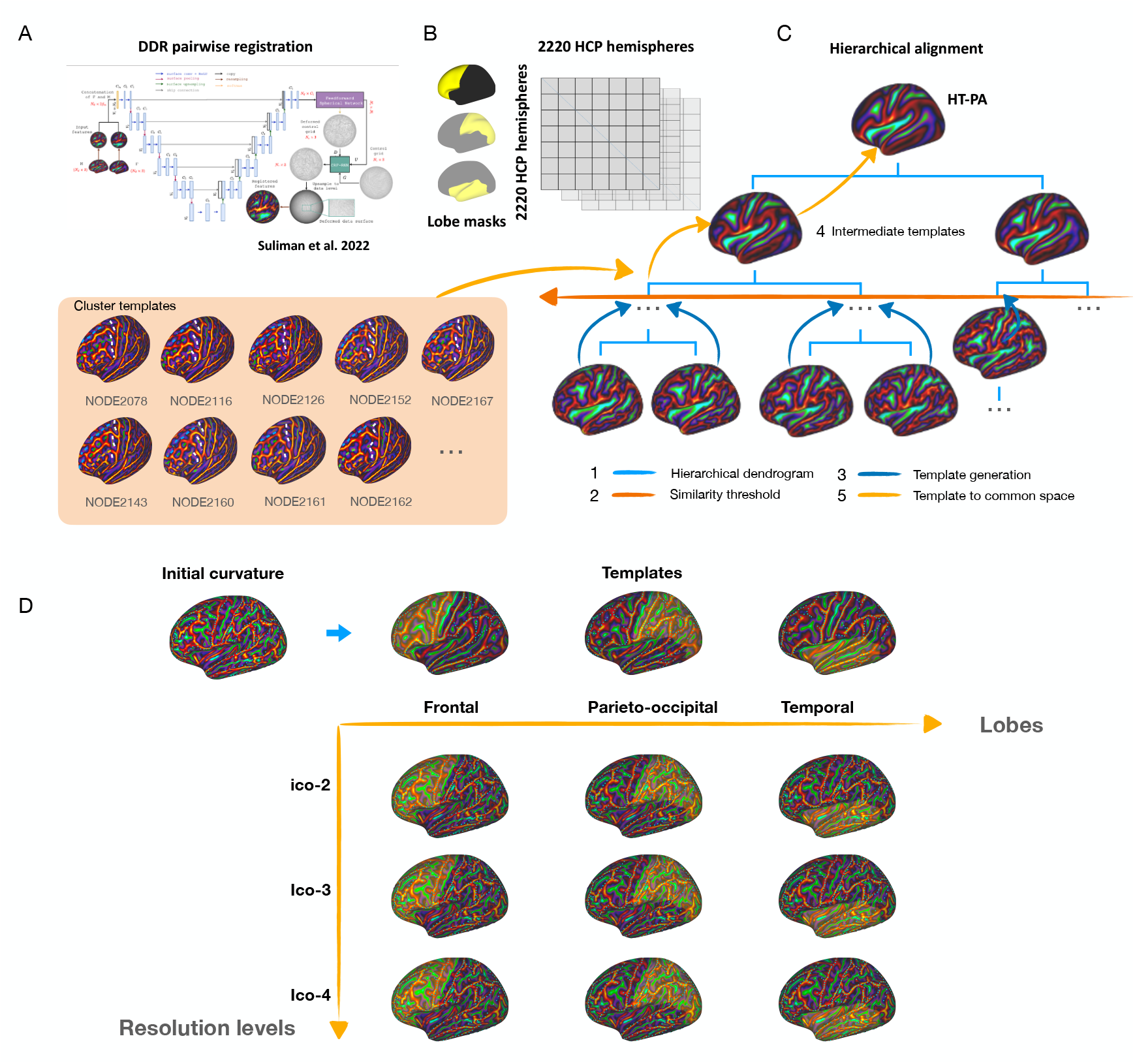
Overview of Multimodal Surface Matching with Hierarchical Templates (MSM-HT). **A**, After mirror-flipping all right hemispheres (into rigid alignment with left hemi-spheres) using Workbench Command, all pairs of sulcal depth maps were co-registered using DDR. **B**, once sulcal depth maps are coarsely aligned, the overlap of cortical folding patterns, for each pair of hemispheres, are separately assessed for each of the frontal, parieto-occipital, and temporal lobes - using a combination of Dice overlap and cross-correlation. **C**, Hierarchical alignment in steps. (1) Agglomerative hierarchical clustering is applied to each of the resulting similarity matrices, (2) the hierarchy is thresholded at the similarity score that returns thirty clusters for each lobe, (3) MSM is then used to co-register curvature maps of individuals within each cluster to generate a template for each cluster - that summarises the shape of folds within that cluster, (4) Intermediate templates were generated at each node of the dendrogram, by aligning pairs of templates until all images were merged into HT-PA at the top. (5) Finally, all templates are further registered through the hierarchical path to a common space. **D**, Combining the registration across three lobes. Mappings are combined across lobes by iterating alignment: first to frontal lobe templates, then parietal and finally temporal. This process repeated through 3 resolution levels of the MSM registration.

#### Optimising pairwise alignment with DDR

DDR is a deep learning-based cortical surface registration algorithm that consists of a rigid-body rotation network and a non-rigid deep-discrete deformation network. In each case, the underlying networks take the form of a U-Net surface convolution neuronal network (CNN), operating on the spherical mesh domain. The rotation network learns to rigidly align all examples to a single population-average template. The non-linear deformation network then learns a non-linear spherical displacement field that aligns an input (or moving) sulcal depth (SD) map with a target (or reference) SD map. The displacement field is optimised for a control grid defined on a lower-resolution icosphere compared to the input and reference feature maps. In this work, the control grid is set to ico-2 (with a resolution of 162 vertices), while the input and reference feature maps are typically at ico-6 (with a resolution of 40,962 vertices). In the original DDR paper the reference map was always the HCP population-average MSMsulc template; in this paper, however, we retrain the deformation network to perform pairwise alignment of individual subject maps instead. We use a mean squared error (MSE) loss to optimise the rotation network and and a combination of MSE and cross-correlation (CC) loss to optimize the deep discrete network. The optimization is performed using the Adam optimizer with a learning rate of 10^*−*4^. Performance was evaluated from 5-fold cross-validation on the HCP dataset through all possible pairwise combinations. More details on the implementation of DDR are expanded in the Supplementary Note Training and validation of DDR and in Suliman et al. (32).

#### Clustering of cortical folding patterns

Following DDR alignment between all pairs of hemispheres, the overlap of cortical folds was assessed from a combination of CC of the entire sulcal depth map and Dice overlap of the gyral crown. Since Dice is normally a measure of segmentation quality, we first thresholded the SD at the 30th percentile (to retain only gyri) and then binarised the result to form a segmentation mask. These similarity metrics were chosen because CC can be seen as being sensitive to the similarity of the general pattern of features. In contrast, Dice is more sensitive to the location of the sulci and gyri. This resulted in a similarity matrix for each lobe, from which individual hemispheres could then be grouped together using agglomerative hierarchical clustering. This establishes a dendrogram which encodes the distance between different surfaces based on the similarity matrix. Initially, each surface stands as its own cluster. In each iteration of the algorithm, clusters are merged in ascending order of distances until a hierarchical path interconnects all data points (see Supplementary Note Evaluation of MSM-HT registration for detailed explanation). We then threshold each dendrogram at the level of 30 clusters (cluster size < 5 not included), where 30 was chosen as it was felt that this represented a manageable number of templates - with sufficient detail to compare against O’Leary et al. (55) - without underpowering individual cluster analyses. Evaluations of alternative approaches to clustering are evaluated in the Supplementary Note Evaluation of MSM-HT registration.

#### Template generation

Once cluster membership was defined, templates were generated through implementing coarse-to-fine alignment with MSM. This optimises warps over a series of control-point grids of increasing resolution - from ico-2 to ico- 4 (2562 vertices) to first perform highly regularised alignment of major folds, before refining alignment of more fine-scale detail. Warps are optimised through discrete optimisation which balances a data similarity (feature overlap) measure, with a regularisation term that penalises non-biologically-plausible deformations (34). Data similarity is implemented as cross-correlation, with regularisation imposed by calculating the hyperelastic strain of the deformation field; this trade-off is balanced with a tunable hyper-parameter *λ*, with all config files used for this paper available from https://github.com/Yrong-Guo/Hierarchical-surface-registration/tree/main/msm_config. To optimise the sharpness of the generated templates, MSM was iterated over multiple rounds. with the first round driven by SD maps, but all subsequent rounds driven to align curvature. This follows the standard protocol for template generation used by (77, 78).

#### Hierarchical alignment to population-average space

While the previous stage is sufficient to co-align hemispheres within each cluster, it presents no guarantees that cortical maps align between clusters. Thus, to enable comparison of features across all clusters, we first generate intermediate templates: following the MSM-HT process up to the top of the dendrogram, through aligning pairs of templates until all images are merged into a full population-average (HT-PA) at the top.

Next, individual hemispheres are brought into HT-PA space, by incrementally aligning each example through the template hierarchy. Since templates are defined separately for each lobe, this requires masking of the cortical feature maps to ensure that only folds from the lobe in question are used to drive alignment to each template. Then mappings are combined across lobes by iterating alignment: first to frontal lobe templates, then parieto-occipital and finally temporal. Since regularization can cause deformation at the margins of the lobar masks, potentially affecting neighbouring regions, its impact is mitigated through iterating template alignments for each of the 3 resolution levels of the MSM registration (Fig. 7D). While this process may seem elaborate, it is necessarily complicated due to the vast heterogeneity of folding patterns and the limited co-occurrence of patterns across lobes. This iterative approach prevents the registration of one lobe from adversely influencing other brain regions, maintaining the integrity of the lobar-specific alignments.

### Lobar folding asymmetry

Lobar asymmetry was assessed by calculating the proportion of left hemisphere examples used to generate each template. Referred to as the Left Hemisphere Rate (LHR) this returned a value between 0 and 1, where 1 indicates that all the hemispheres in a cluster are left hemispheres. Clusters were categorized as leftward-or rightward-asymmetric based on whether their LHR was greater than 0.5. We used Mann-Whitney U Test to determine significance of the difference between each pair of lobes, followed by Bonferroni correction for multiple comparisons. On observation of significantly greater asymmetry of the temporal lobe, we characterised the differences through manual labelling and quantification of the rate of branches and interruptions in each template. We also estimated the total surface area of this region (defined by Desikan-Killiany atlas), normalised by the total ipsilateral surface area. The significance of this difference was evaluated using a two-sample t-test.

### Generalisation on developmental cortical folding

For investigating cortical folding variants in neonates, we replicated the MSM-HT framework (described in Hierarchical MSM registration (MSM-HT)) in the dHCP and BIBS cohorts. Since neonatal cortical feature maps display very different imaging properties, the DDR deformation network was retrained on the dHCP dataset, utilising the same architecture and optimization procedures with the same parameters as the HCP. Lobar masks were generated from the DRAW-EM neonatal atlas (70–72). MSM-HT was then run and evaluated in exactly the same way as for the HCP-YA data. Finally, neonatal and adult templates were paired to assess the similarity of each pair in the HT-PA space (assessed from cross-correlation).

### Heritability of cortical folding

The role of genetics on higher-order folding was assessed by comparing the cluster membership of monozygotic (MZ) and dizygotic (DZ) twins, using only the HCP dataset. A Chi-square test was conducted to assess whether there is a significant association between the twin type and their likelihood of being in the same cluster. The p-value was corrected using an FDR of 0.05 for multiple comparisons between lobes and hemispheres. The Odds Ratio (OR) was also calculated to measure the magnitude (effect size) of the association. An OR > 1 suggests that MZ twins are more likely to present in the same folding cluster than DZ twins.

### Folding-function relationship

We hypothesised that a more accurate alignment of cortical folds could improve the alignment of functional activations, since the development of functional regions is known to be partially related to the development of cortical folds. If true this should also mean that individuals within the same cluster should share topologically similar cortical areal topographies (as estimated from the HCP multimodal cortical areal parcellation). Thus, to probe the spatial correspondence between structure and function, the MSM-HT and MSM-P warps were used to map tfMRI, and each individual’s HCP_MMP1.0 parcellation into the HCP-HT and HCP-P folding template spaces.

For tfMRI, we used specific contrasts from the Language (Story, Math), Working Memory (N-back tasks), Social Cognition (Theory of Mind), and Motor (movement of left/right hand, left/right foot, tongue) tasks. These contrasts were chosen for their well-documented spatial activations. The group-average tfMRI activation was calculated using a GLM-based fMRI analysis with mixed effects FLAMEO (67). *Cluster mass* was used to measure the strength and spatial concentration of significantly activated vertices (as defined in (34)), where higher values indicate better overlap of tfMRI activations across the population under consideration. The significance of the difference between MSM-HT and MSM-P were evaluated using two-sample t-tests, followed by FDR of 0.05 for multiple comparisons.

Individual subject HCP_MMP1.0 parcellations were downloaded from the BALSA database. These define cortical areas from a combination of multi-modal areal features including myelin maps, cortical thickness, tfMRI contrast maps, resting-state fMRI functional connectivity (FC) maps, and continuous, topographically organised gradients of resting-state fMRI functional connectivity (44). Thus these represent a more comprehensive measure of cortical organisation than tfMRI alone. Overlap of areal features across individuals after alignment was assessed by the areal uncertainty of each vertex: calculated as one minus the highest proportion of individuals sharing the same label at that location. To ensure that the areal uncertainty was not inflated by large cluster sizes, we calculated it both on the original clusters and after breaking the clusters into comparable sizes based on the dendrogram.

## ACKNOWLEDGEMENTS

The authors would like to acknowledge the participants in the HCP, dHCP and BIBS. We thank Matthew F. Glasser and David C. Van Essen for their invaluable input during the design of this study. We also thank Gareth Ball and Katherine R. Long for their insightful feedback. HCP-YA data were provided by the Human Connectome Project, WU-Minn Consortium (Principal Investigators: David Van Essen and Kamil Ugurbil; 1U54MH091657) funded by the 16 NIH Institutes and Centers that support the NIH Blueprint for Neuroscience Research; and by the McDonnell Center for Systems Neuroscience at Washington University in St. Louis. The HCP was approved by the internal review board of Washington University in St. Louis (IRB #201204036). The dHCP project was funded by the European Research Council (ERC) under the European Union Seventh Framework Programme (FR/2007-2013)/ERC grant agreement no. 319,456. BIBS data was funded by EU-AIMS (European Autism Interventions)/EU AIMS-2-TRIALS, an Innovative Medicines Initiative Joint Undertaking under Grant Agreement No. 777394. The School of Biomedical Engineering and Imaging Sciences is supported by the Wellcome EPSRC Centre for Medical Engineering at King’s College London (WT 203148/Z/16/Z) and the Department of Health via the National Institute for Health Research (NIHR) comprehensive Biomedical Research Centre (BRC) award to Guy’s & St Thomas’ NHS Foundation Trust in partnership with King’s College London and King’s College Hospital NHS Foundation Trust. We acknowledge infrastructure support from the NIHR Mental Health BRC at South London and Maudsley NHS Foundation Trust, King’s College London. The views expressed are those of the author(s) and not necessarily those of the NHS, the NIHR or the Department of Health and Social Care. For the purposes of open access, the authors have applied a Creative Commons Attribution (CC BY) licence to any Accepted Author Manuscript version arising, in accordance with King’s College London’s Rights Retention policy. We acknowledge the support by the European Research Council under the European Union Seventh Framework Programme (FP/2007–2013)/ERC Grant Agreement No. 319456.

Y.G. was supported by the King’s-China Scholarship Council PhD Scholarship. E.C.R is supported by funding from the MRC methodology grant (MR/V03832X/1). E.C.R. and J.O. received support from the Medical Research Council Centre for Neurodevelopmental Disorders, King’s College London (MR/N026063/1). J.O. is supported by a Sir Henry Dale Fellowship jointly funded by the Wellcome Trust and the Royal Society (206675/Z/17/Z). V.K. is supported by an MRC Transition Support Award (MR/V036874/1). The authors acknowledge use of the King’s Computational Research, Engineering and Technology Environment (CREATE) (https://doi.org/10.18742/rnvf-m076).

