## Supplementary Information for "Motifs of brain cortical folding from birth to adulthood: structural asymmetry and folding-functional links"

### Supplementary Figures

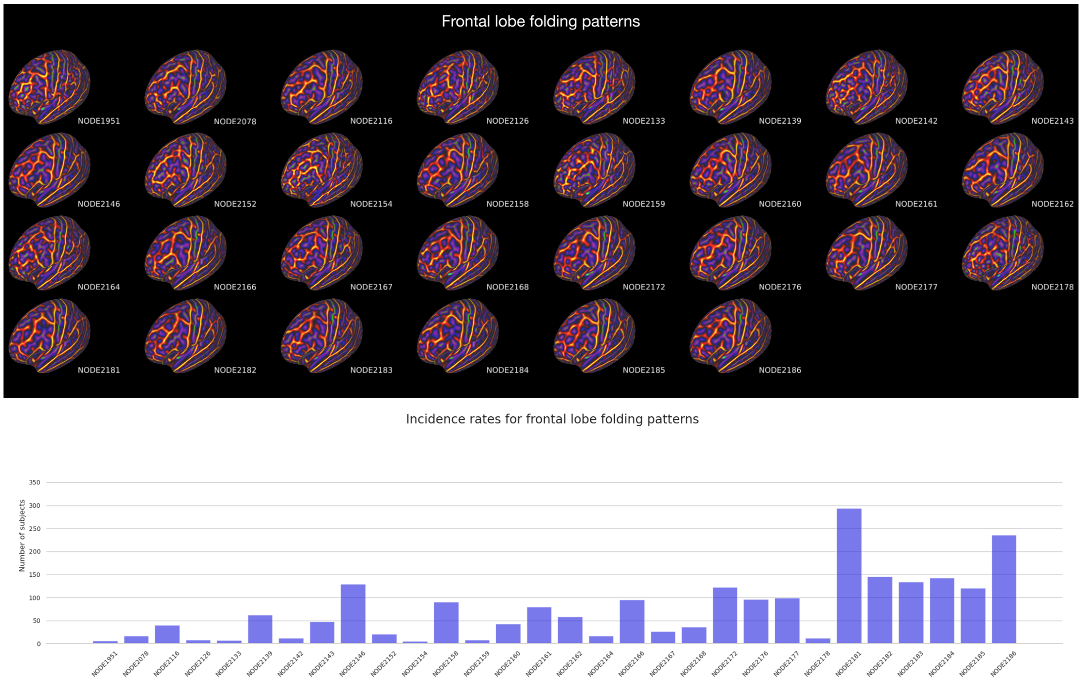


**Fig. 1.** Cortical folding patterns and incidence rates of the frontal lobe.

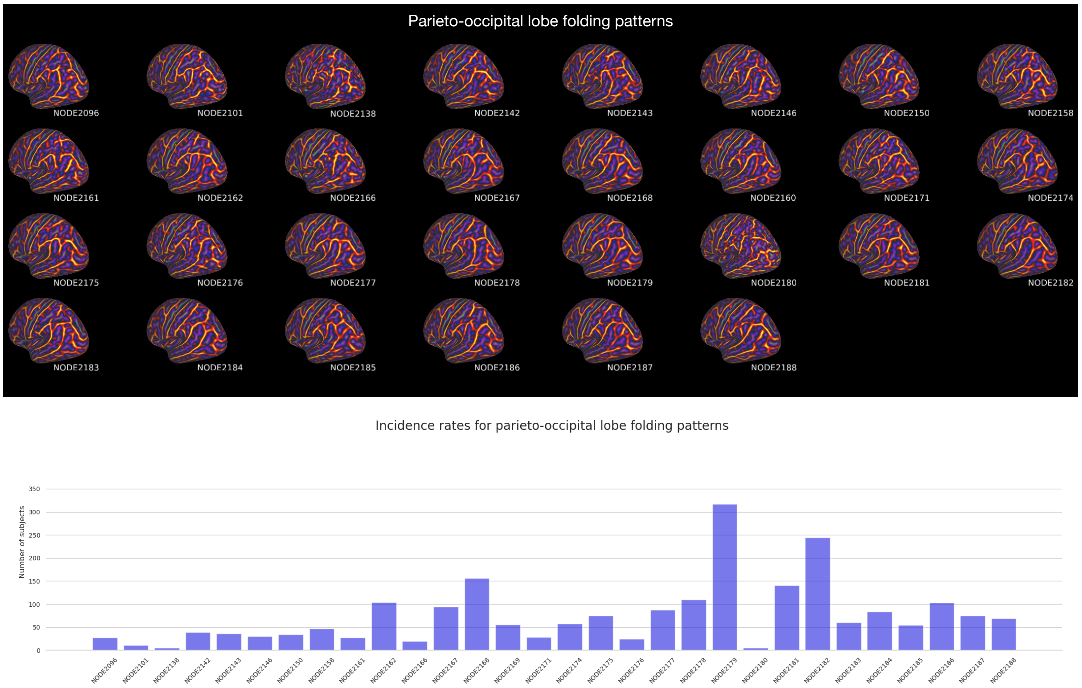


**Fig. 2.** Cortical folding patterns and incidence rates of the parieto-occipital lobe.

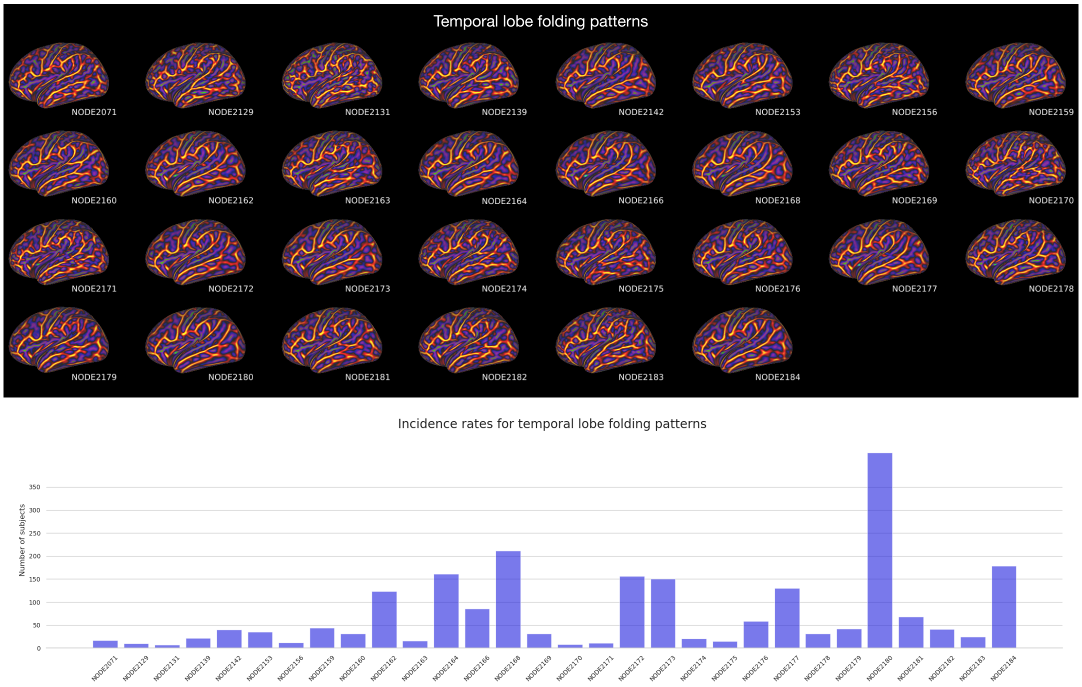


**Fig. 3.** Cortical folding patterns and incidence rates of the temporal lobe.

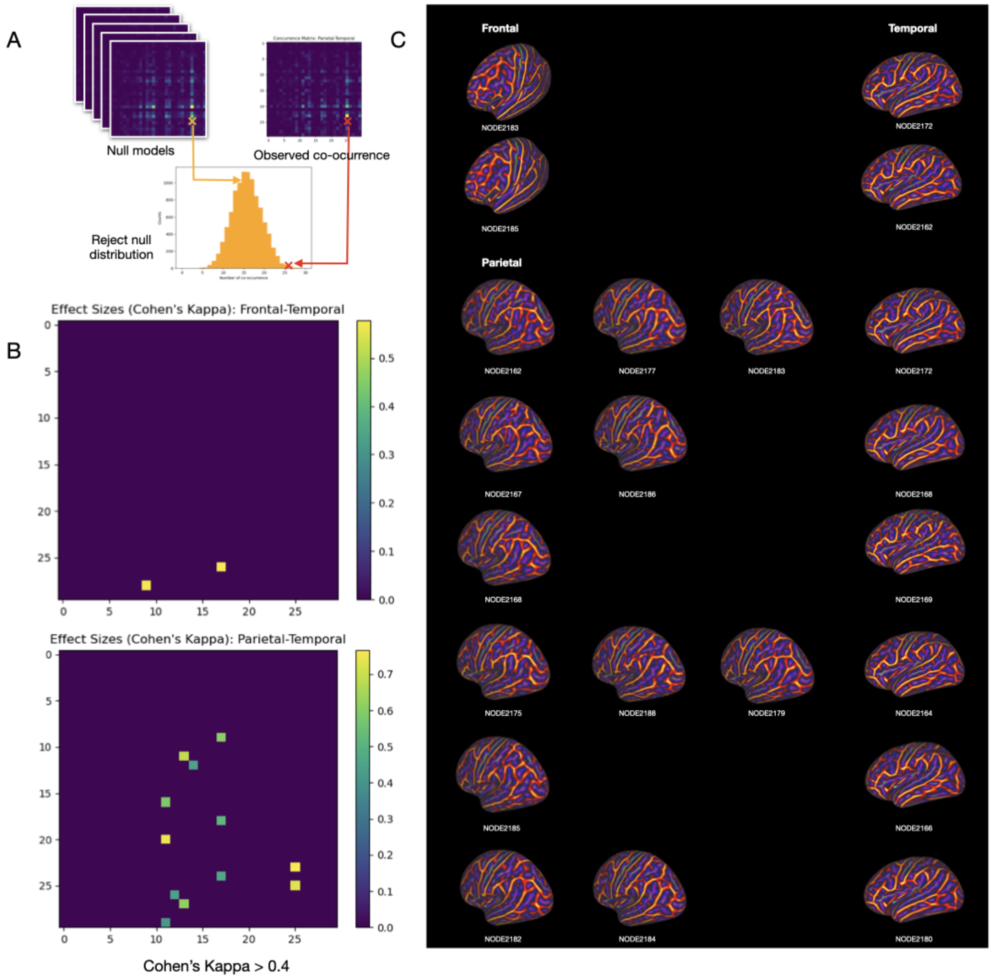


**Fig. 4.** Co-occurrence between lobes. **a.** Construction of null co-occurrence matrices by permutation of subjects across different clusters. 0.05 is settled as a threshold to reject null models. **b.** co-occurrence with Cohen's $\kappa$ > 0.4. **c.** The pairs of cortical folding patterns that significantly co-occur (p-corrected < 0.05, $\kappa$ > 0.4 ) between lobes organised by row. There are co-occurrences of folding across the entire brain with 13 frontal-temporal, 46 parietal-temporal, and 1 frontal-temporal co-occurrences reaching significance; however, effect sizes are generally small $\kappa$ < 0.2. In total 14 pairs (12 parietal-temporal and 2 frontal-temporal) achieve $\kappa$ > 0.4 indicating moderate to substantial agreement; however, these are largely characterised by larger cluster sizes and more common folding patterns.

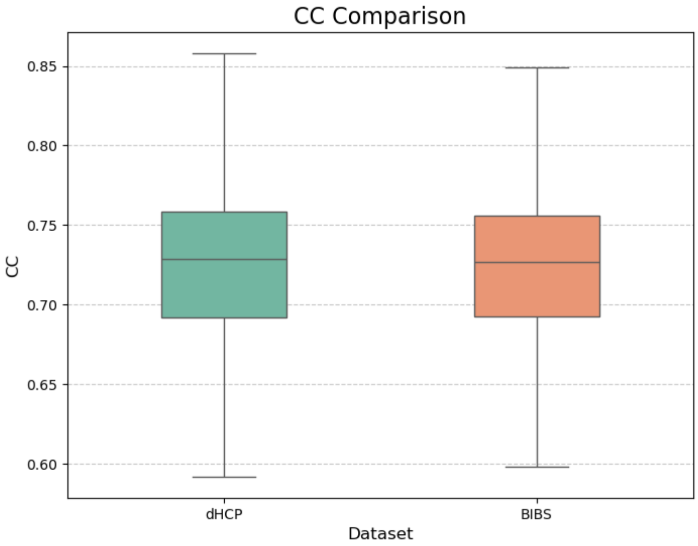


**Fig. 5.** The CC accuracy values comparing dHCP (training) and BIBS (testing) dataset. Results of this evaluation indicated that the mean CC values for the dHCP and BIBS datasets were 0.717 and 0.720, respectively; the difference in distributions was significant ($p = 6.44e - 10$) but the effect size was small (Cohen's d = -0.044). These values suggest DDR performs consistently across training and testing datasets.

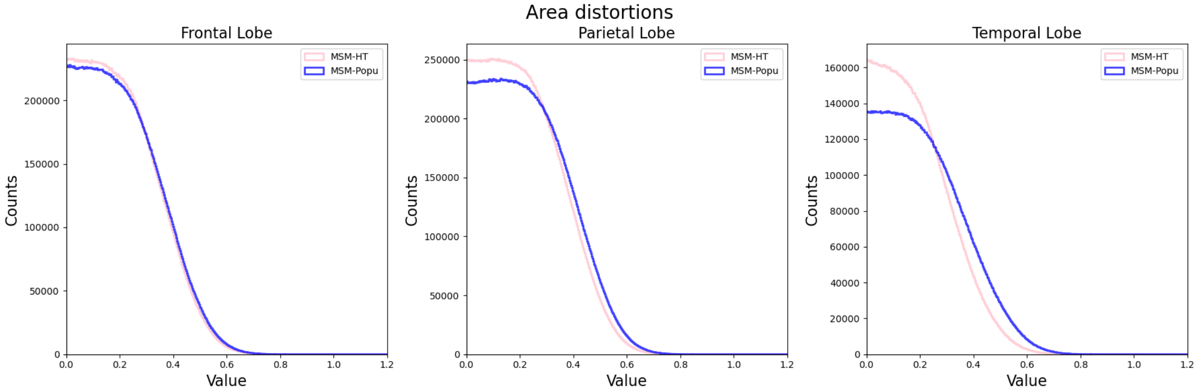


**Fig. 6.** The area distortion comparing MSM-HT and MSM-Popu

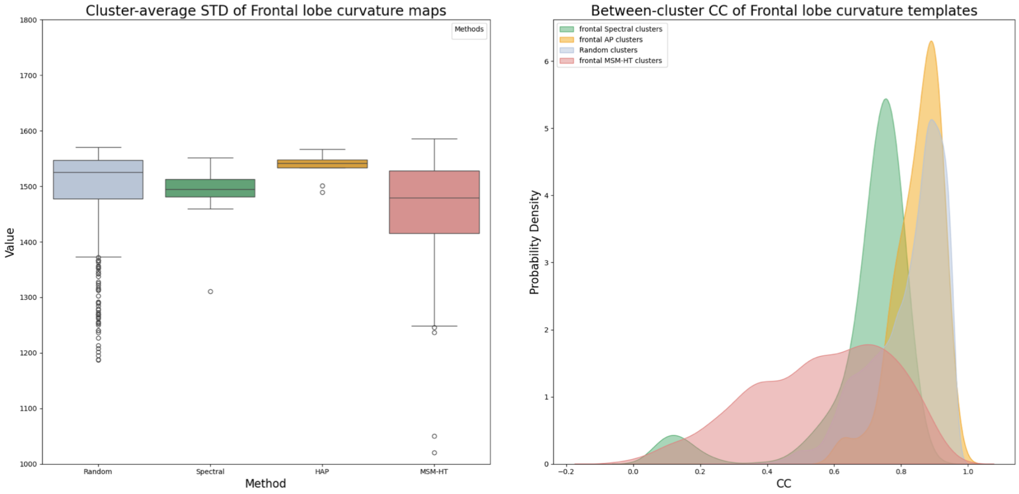


**Fig. 7.** Within- and between-clusters performance. **a.** Within-cluster STD and between cluster CC compared between four methods: 1) Random generated clusters, 2) Spectral clustering (SP), 3) Hierarchical affinity propagation (hierarchical AP), and 4)MSM-HT. Quantitatively, the STD map was found to be significantly lower in MSM-HT clusters, followed by SP, hierarchical AP, and random clusters, indicating less registration uncertainty within MSM-HT clusters ($p<0.05$). **b.** The between-template similarity shown by CC for MSM-HT clusters is higher than SP, hierarchical AP and random clusters, indicating more compact and well-defined clusters.

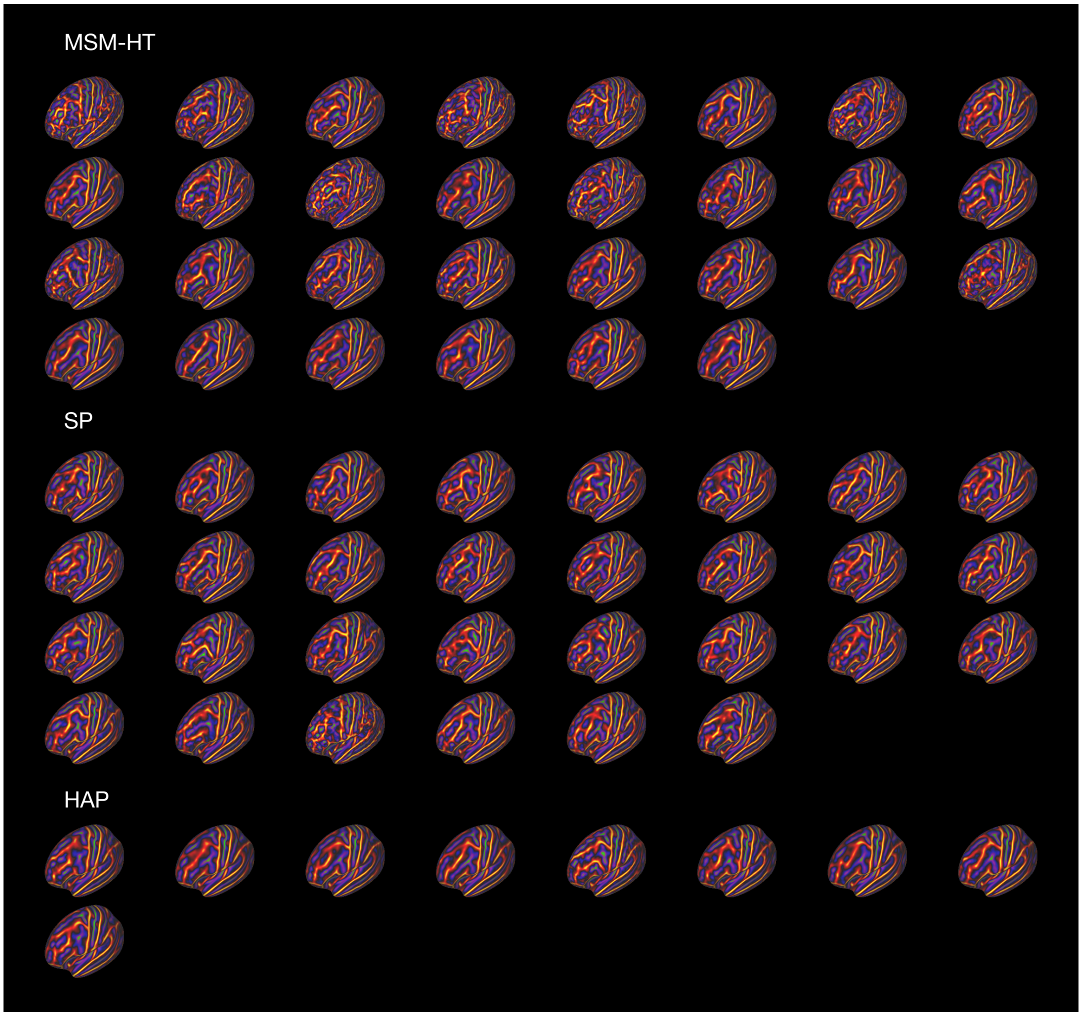


**Fig. 8.** Visual comparison between MSM-HT, SP, and hierarchical AP. MSM-HT generates visually sharper mean maps with more detail and greater variability than templates generated from SP, hierarchical AP, or random allocation. For example, single-segment precentral sulcus clusters such as clusters 3 and 14 are only uncovered by MSM-HT, but only two or three segments were found in AP clusters, and only two segments of precentral sulcus are found in hierarchical AP clusters. An example of this is clusters 1 and 6 in MSM-HT against cluster 17 in SP clusters and cluster 4 in AP clusters. Random clusters, which serve as a baseline for comparison, exhibited the least defined patterns among all methods evaluated. These clusters generally lack details of segments or branching patterns and often resemble the overall population average.

#### Supplementary Notes

**Deep learning-based neonatal pipeline** All dHCP and BIBS data are processed using the dHCP deep learning-based neonatal pipeline, detailed in (1). In brief, T2w images are first bias-corrected with the N4 algorithm (2), then rigidly and affinely aligned to the 40-week neonatal volumetric template (3). Then a 3D CNN with U-Net architecture learns to predict white matter and pial surfaces by deforming the dHCP 40-week population-average surface template, until it fits the inner and outer cortical boundaries of each subject's T2w MRI. Anatomical surfaces were then inflated using a GPU parallelised implementation of Freesurfer's *mris_inflate* (4, 5). Finally, an unsupervised learning-based spherical mapping approach was used to generate spheres for each individual, by deforming the 40-week template sphere (generated from FreeSurfer's *mris_sphere*) to minimise metric distortions between the learnt sphere and each individual's white matter surface. This results in a comprehensive set of white, pial, inflated and spherical surfaces for each subject, with corresponding metric files summarising cortical thickness, curvature and sulcal depth. However, the pipeline does not run segmentation; therefore we define anatomical lobes from the DRAW-EM (6) segmentations integrated into original dHCP pipeline (7). These labels, generated in volume space, were projected to the surface using ConnectomeDB *volume-label-to-surface-mapping*.

**Training and validation of DDR** For HCP-YA, DDR was trained on 1110 subjects in a five-fold cross-validation manner. Alignment accuracy in the test cohort was evaluated using mean square error (MSE) and cross-correlation (CC). The five-fold average CC is 0.750.

For the neonates, DDR was trained on 717 dHCP subjects with five-folds and evaluated in 100 BIBS subjects. The image acquisition and cortical processing was identical for both cohorts. Results of this evaluation indicated that the mean CC values for the dHCP and BIBS datasets were 0.717 and 0.720, respectively, suggesting DDR performs consistently across training and testing datasets (Supplementary Fig.5).

**Evaluation of MSM-HT registration** To validate the MSM-HT pipeline, we conducted several validation steps to benchmark its performance against conventional methods.

**Validating registration performance** We generate metric maps that convey the mean and standard deviation (STD) of the curvature for each vertex across all individuals within each cluster, aligned by MSM-HT and MSM-P registration. Paired examples were compared for the STD values with a Wilcoxon signed-rank test. The result shows that MSM-HT is visually sharper (see main paper), and significantly reduced the variance across subjects within the clusters ($p < 0.001$). Furthermore, after combining the registration across different lobes, the STD values are still significantly lower within each cluster than MSM-P ($p < 0.001$).

Areal distortion for each vertex is defined as the average of the distortion for all triangles adjacent to the vertex $p$, $1/N_p \sum_i log_2(AD_i/AO_i)$, where $AO_i$ and $AD_i$ are the area of triangle $i$ before and after distorted and $N_p$ is the number of triangle adjacent to vertex $p$. To ensure a fair comparison, we controlled for area distortion - adjusting regularisation terms (i.e. lambda, see main paper) until the 95th percentile of area distortions matched across both datasets (Distribution of the area distortion shown in Supplementary Fig.6).

**Justification of clustering approach** In agglomerative hierarchical clustering, the distance between two clusters was determined by the maximum distance between data points (known as *complete linkage*) to generate more compact clusters. At the point where the resultant hierarchy was thresholded, thirty cortical folding clusters were revealed (clusters with sizes smaller than five were excluded to ensure the clusters were representative).

Many different clustering approaches exist, including spectral clustering (SP) and hierarchical affinity propagation (Hierarchical AP - used by (8, 9)). Therefore, we validate the chosen approach against these methods in terms of: a) the sharpness of the generated templates (validated from within template curvature STD); b) the diversity of the generated templates - assessed from the CC of different templates derived from the same method. This is compared against a baseline generated by randomly allocating individual hemispheres to 500 clusters (with cluster sizes scaled to match the distribution of the MSM-HT templates). The implementation of Both SP and AP are built-in functions of *sklearn.cluster*, and hierarchical AP follows the detailed description in Duan et al. (9). In order to save on computation costs, this is only performed in the frontal lobe. The cluster curvature STD and between-cluster CC are shown in Supplementary Fig. 7, and the visual comparison between MSM-HT, SP, and hierarchical AP are shown in Supplementary Fig. 8.

#### Supplementary Results

**Co-occurrence analysis of cortical folding patterns across lobes** To test the significance of the co-occurrence, we generated 10000 null models that were built by randomly permuting the cluster memberships of the subjects. This approach creates a distribution of co-occurrence expected by chance, allowing us to calculate the p-values of each pair of folding templates (corrected using the FDR method to control for multiple comparisons). To ensure meaningful significance, we evaluated the effect size of co-occurrence using Cohen's Kappa ($\kappa$). We focused on $\kappa > 0.4$ to target co-occurrences that fall into the moderate to substantial agreement range.

Results, shown in 4, point to the co-occurrence of folding across the entire brain with 13 frontal-temporal, 46 parietal-temporal, and 1 frontal-temporal co-occurrences reaching significance ($p - corrected < 0.005$); however, effect sizes are generally small $\kappa$ < 0.2. In total 14 pairs (12 parietal-temporal and 2 frontal-temporal) achieve $\kappa$ > 0.4 indicating moderate to substantial agreement; however, these are largely characterised by larger cluster sizes and more common folding patterns.

### Supplementary Discussion

**Co-occurrence analysis of cortical folding patterns across lobes** Although genetic influences on cortical surface area (CSA) are region-specific (10), the development of cortical regions is highly interconnected. Previous studies of the structural covariance network start from the covariance of cortical thickness, surface area, and significances were found (11, 12). Our study further extends the focus into cortical folding patterns, where parieto-occipital folding patterns seem to co-occur more with temporal lobe patterns. However, since the significant pairs are characterised by larger cluster sizes and more common folding patterns, this might suggest that the analysis is underpowered due to relatively insufficient data.

Structural covariance networks, complemented with structural and functional connectivity, have offered valuable insights into how different functional regions coordinate. Lerch et al. (13) first looked into the correlation between cortical thickness in different parts of the brain and found covariance between different language areas, which agrees with the connection of white matter tracts. Recently, genetic factors have been found to play a crucial role in structural covariance of cortical thickness (14), which is related to the functional gradients and conserved in primates. Our method provides a means for calculating the co-occurrence of cortical folding patterns between different lobes. The significant co-occurrence of folding patterns between the temporal and parietal lobes aligns with existing literature suggesting close anatomical relationships between the temporal and parietal lobes, which may contribute to integrated processing tasks such as language comprehension and spatial awareness. However, it is important to consider the potential methodological bias due to cluster size. While the permutation tests were designed to account for cluster size by shuffling memberships while maintaining cluster size, this inherent bias cannot be fully ruled out. Further investigation is needed to determine whether these interactions are biologically meaningful or primarily an artefact of cluster size.
